# Lipid droplet and peroxisome biogenesis occur at the same ER subdomains

**DOI:** 10.1101/188433

**Authors:** Amit S. Joshi, Vineet Choudhary, Tim P. Levine, William A. Prinz

## Abstract

Nascent lipid droplet (LD) formation occurs in the ER membrane^1^^-^^4^. It is not known whether LD biogenesis occurs stochastically in the ER or at subdomains with unique protein and lipid composition. We previously identified ER subdomains in *S. cerevisiae* that contain Pex30, a reticulon-like ER-resident membrane protein^5^. There are ~25 Pex30-containing puncta in the ER per cell. These sites are regions where preperoxisomal vesicles (PPVs) are generated^5^. Here we show that Pex30 subdomains are also the location where most nascent LDs form. Mature LDs remain associated with Pex30 subdomains and the same Pex30 subdomain can simultaneously associate with a LD and a PPV. Pex30 subdomains become highly enriched in diacylglycerol (DAG) during LD biogenesis, indicating they have a unique lipid composition. We find that in higher eukaryotes multiple C2 domain containing transmembrane protein (MCTP2) is the functional homologue of Pex30; MCTP2 resides in ER subdomains where most nascent LD biogenesis occurs and that are often associated with peroxisomes. Together, these findings indicate that most LDs and PPVs form and remain associated with conserved ER subdomains and suggest a link between LD and peroxisome biogenesis.

---

LDs are composed of a core of neutral lipids, triacylglycerides (TAG) and sterol esters, covered by a phospholipid monolayer containing proteins. LD biogenesis probably begins when neutral lipids in the ER coalesce to form lens-like structures between the leaflets of the membrane^6^. These lenses subsequently grow larger and bud from the ER, forming mature LDs^7^^-^^10^. How locations of LD biogenesis in the ER membrane are determined is not known but there is evidence that biogenesis might occur at specialized sites^1,4^.

We wondered whether LD biogenesis occurs at Pex30 domains in the ER for two reasons. First, there are about 10-fold more Pex30 domains than there are PPVs in cells^5^, suggesting these domains have other functions. Second, recent evidence suggests that some proteins play dual roles in the biogenesis of both LDs and peroxisomes; the Kapito group found that two proteins required for peroxisome biogenesis, Pex3 and Pex19, insert membrane-embedded proteins into the surface of LDs at ER subdomains^11^.

To determine whether nascent LDs mature at Pex30 subdomains, we visualized LD biogenesis in a *S. cerevisiae* strain in which LD formation can be controlled. Four enzymes produce neutral lipids in this yeast: Are1 and Are2, which generate steryl esters, and Lro1 and Dga1, which synthesize TAG. Cells lacking all four proteins lack neutral lipids and LDs. We used a strain in which the galactose regulatable promoter *GAL1* controls expression of *LRO1* and the other three neutral lipid-synthesizing enzymes are not produced (*GAL1-LRO1* 3Δ). When this strain is grown in a medium containing raffinose, it lacks LDs but begins to produce them when galactose is added^12^. The strain also expressed Pex30-2xmCherry and Erg6-GFP, a LD marker. Before *LRO1* induction, Erg6-GFP is on the ER but it localizes to LDs after galactose addition (Fig. 1A). About 70% of Erg6-GFP punctae colocalize with Pex30-2xmCherry (Fig. 1B). Similar results were obtained when nascent LDs were visualized with the lipophilic dye BODIPY (Supplementary Fig. 1A).

**Figure 1.**
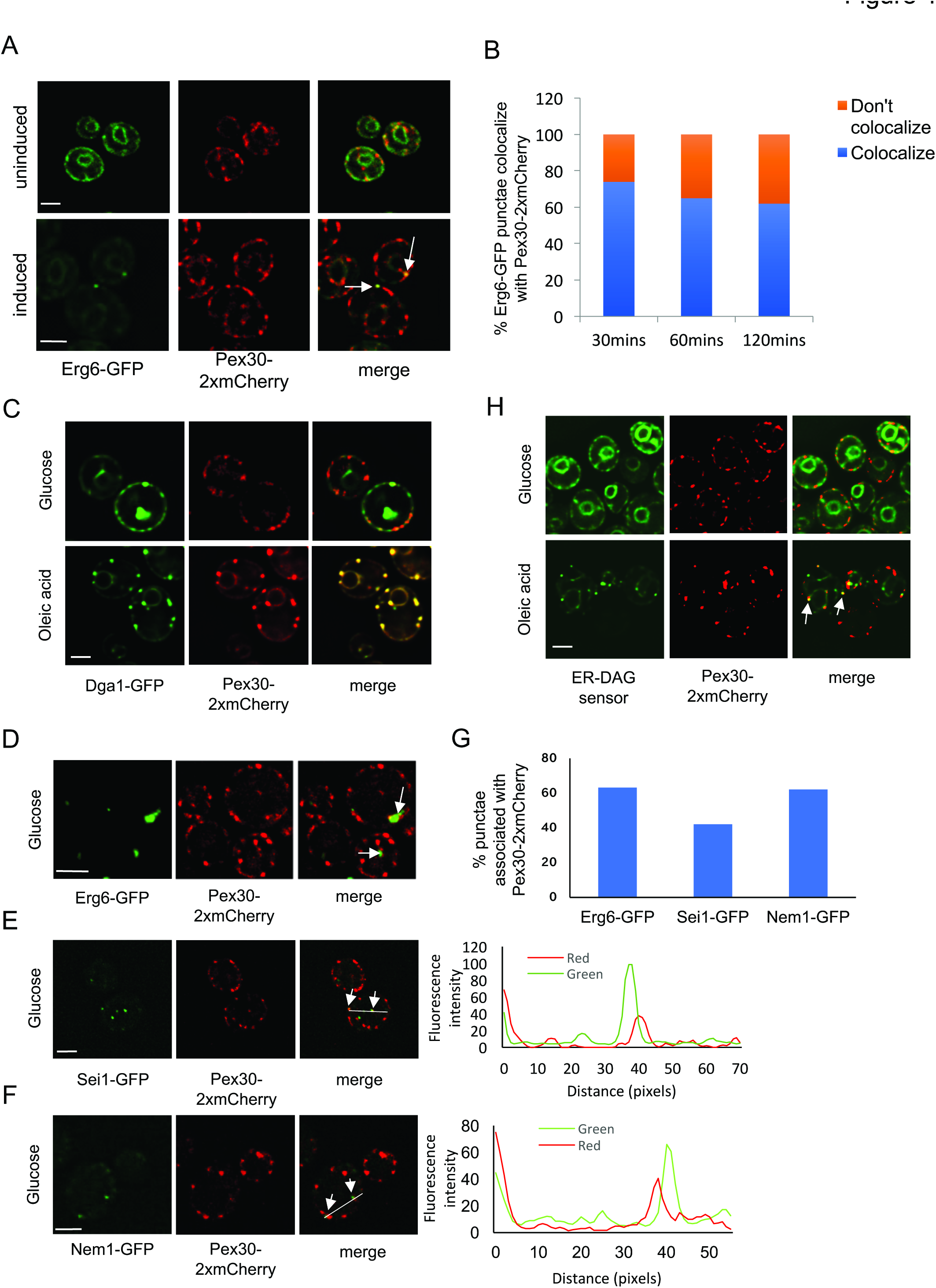
Pex30-containing ER subdomains associate with LDs. **A)** *GAL1-LRO1* 3Δ cells expressing endogenously tagged Erg6-GFP and Pex30- 2xmCherry were visualized growing in a medium containing raffinose (uniduced) or 30 minutes after galactose addition (induced). White arrows indicate colocalization of Erg6-GFP and Pex30-2xmCherry. **B)** Percent of Erg6-GFP punctae that colocalize with Pex30-2xmCherry after galactose addition. **C)** Wild-type cells expressing endogenously tagged Pex30-2xmCherry and the plasmid Yep181-Dga1-GFP were grown to stationary growth phase in SC medium (glucose). The cells were washed, incubated with fresh SC containing 1mM oleic acid, and imaged after 1 hour (oleic acid). **D-F)** Wild-type cells expressing endogenously tagged Pex30-2xmCherry and Erg6-GFP (D), Sei1-GFP (E), or Nem1-GFP (F) were grown in SC to early stationary growth phase phase and visualized live. Arrows indicate sites where Pex30-2xmCherry associates with other makers. Graphs to right of (E) and (F) show signal intensity on white line. **G)** Quantification of percent Pex30-2xmCherry punctae associated with Erg6-GFP, Sei1-GFP, Nem1-GFP; n = at least 300 cells. **H)** Wild-type cells expressing endogenously tagged Pex30-2xmCherry and the ER-DAG sensor were grown as in C. Bars = 3μm. In D,E,F, and H stacks of 3 images with a step size of 0.25μm were taken and deconvolved; images from a single plane are shown.

To verify that most nascent LDs mature at Pex30 subdomains, we induced LD production with a second method. When oleic acid is added to growing cells, they rapidly begin to produce new LDs. We added oleic acid to cells expressing Dga1-GFP from a high copy plasmid and endogenously expressed Pex30-2xmCherry. In growing cells, Dga1-GFP is in the ER but it relocates to the surface of LDs when LD production is induced. We found that Dga1-GFP puncta colocalize with Pex30-2xmCherry within 1-2 hours after oleic addition (Fig. 1C). Lro1-GFP similarly accumulates at sites containing Pex30-2xmCherry after oleic acid addition. (Supplementary Fig. 1B). Together, these findings indicate that most nascent LDs colocalize with Pex30 subdomains in the ER.

Next, we determined whether mature LDs, like nascent LDs, remain associated with Pex30 domains in the ER. Wild-type cells growing in standard growth media contain mostly mature LDs. Therefore, to determine whether Pex30 domains associate with mature LDs, we imaged cells expressing Pex30-2xmCherry and Erg6-GFP. About ~70% of Erg6-GFP puncta were closely associated with Pex30-2xmCherry (Fig. 1D and 1G), suggesting that not only nascent LDs but also mature LDs remain associated with Pex30 subdomains. We also confirmed the presence of Pex30 at ER subdomains associated with LDs by examining the colocalization of Pex30-2xmCherry with other proteins known to be at these regions. Nem1 is part of a phosphatase complex that regulates Pah1, the yeast homologue of lipin, which plays a central role in controlling TAG levels in cells^13^^-^^15^. Nem1 forms puncta on the ER that are near sites of LD biogenesis^16^^,^^17^. We found that ~70% of Nem1-GFP punctae co-localized with Pex30-2xmCherry (Fig. 1E and 1G), consistent with the idea that Pex30 subdomains remain associated with growing and mature LDs. We also determined whether Sei1, the yeast homologue of seipin, is associated with Pex30 subdomains. Seipin plays an important but still poorly understood role in LD biogenesis and it has been found that some Sei1 puncta localize at ER-LD contacts^17^^,^^18^. We found that 40% of Sei1-GFP puncta were associated with Pex30-2xmCherry domains (Fig. 1F and 1G). Consistent with this finding, it was reported that Pex30 co-immunoprecipitates with Sei1^19^. It may be that the Sei1 puncta away from Pex30 puncta are those not associated with LDs. Taken together, these findings indicate that Pex30 subdomains frequently remain associated with mature LDs.

If Pex30 subdomains are indeed sites of LD biogenesis, we speculated that they might become enriched in neutral lipid precursors when nascent LD production is induced. DAG is a TAG precursor used by both TAG-synthesizing enzymes in yeast. To determine whether DAG becomes enriched at Pex30 sites, we developed a sensor to investigate the distribution of DAG in the ER. The DAG-binding tandem C1 domains of Protein Kinase D have been used to sense DAG in membranes^20^. We fused these domains to GFP and the transmembrane domain of Ubc6, a tail-anchored ER protein (ER-DAG sensor). After confirming that the sensor colocalizes with the ER marker RFP-HDEL (Supplementary Fig. 1C), we expressed the sensor in cells expressing Pex30-2xmCherry. When the cells were grow in regular media, the sensor was all over the ER but it became highly enriched in portions of the ER within 1-2 hours after oleic acid addition, forming bright puncta. About 65% of the puncta colocalized with Pex30-2xmCherry (Fig 1H). This finding indicates that some Pex30 subdomains become highly enriched in DAG when LD formation is induced, consistent with the idea that the subdomains are regions where LDs form. Why DAG does not become enriched at all Pex30 subdomains is unclear but suggests that these subdomains have multiple functions Interestingly, we found that the soluble C-terminal portion of the Pex30 (250-523aa) binds DAG immobilized on membranes (Supplementary Fig. 1D), suggesting that Pex30 could be regulated by DAG accumulation at sites of LD biogenesis.

To investigate the role of Pex30p in LD biogenesis, we examined LDs in cells lacking Pex30 (*pex30*Δ). LDs in these cells were often more clustered and smaller than those in wild-type cells (Fig. 2A-F). The decrease in LD size is probably not caused by a change in the level of neutral lipids in *pex30*Δ cells (Fig 2G and H). Although these cells had a small but significant decrease in TAG levels, this change is probably not large enough to decrease LD size. The change in LD size in *pex30*Δ cells could be because Pex30 affects membrane tension at sites of LD biogenesis. A similar role has been proposed for REEP1, a mammalian reticulon-like ER-shaping protein ^21^ ^22^.

**Figure 2.**
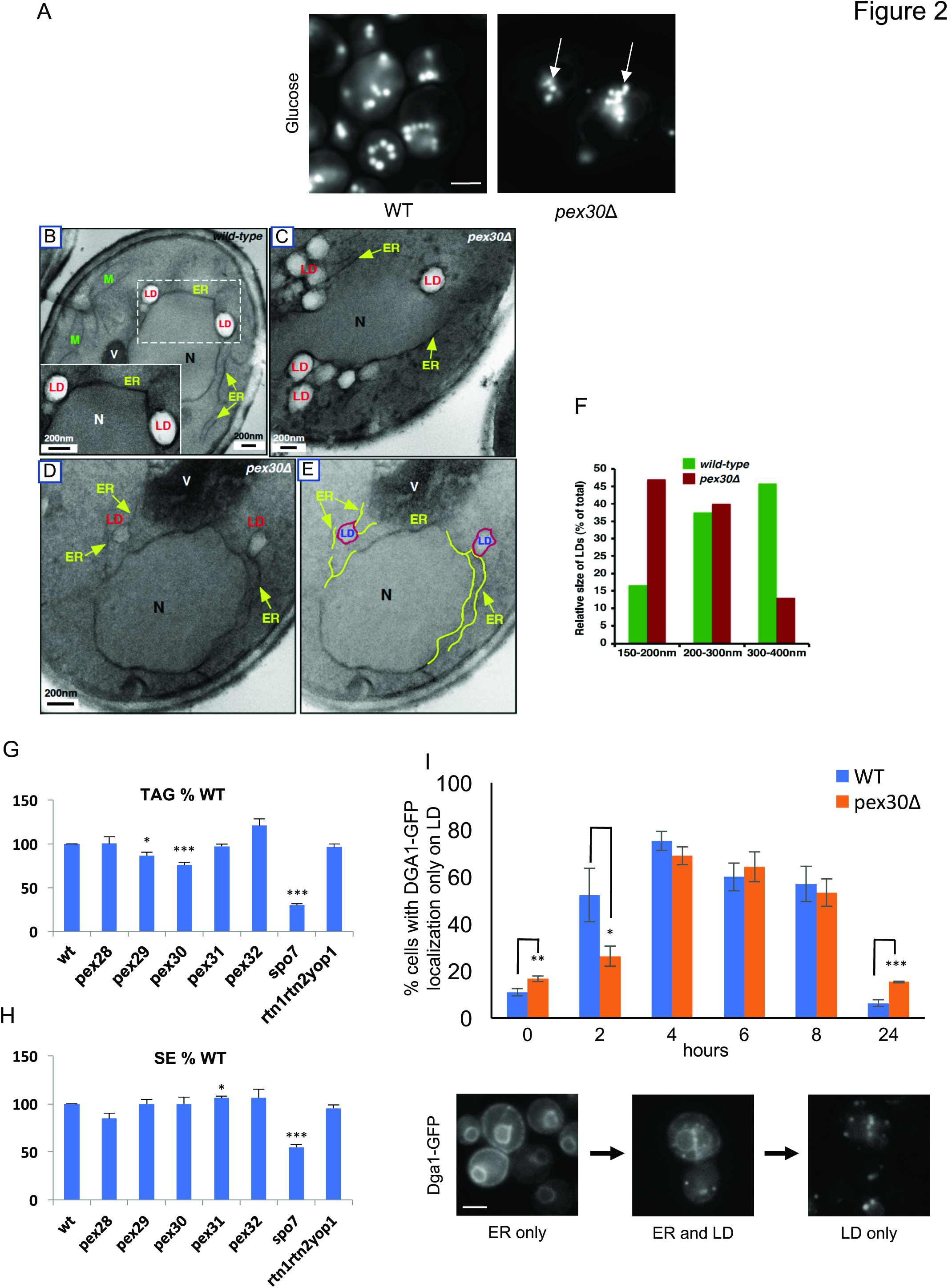
Cells lacking Pex30 have altered LDs. **A)** Cells in SC were grown to early stationary growth phase and stained with BODIPY to visualize LDs. Stacks of 10 images with a step size of 0.2μm were taken; images from a single plane are shown. Arrows indicate clustered LDs. Scale bar 3μm. **B-E)** Cells growing in SC were fixed and visualized by EM. Yellow arrows indicate ER. V; vacuole, N; Nucleus, LD; Lipid droplets. E shows the same image as D but with portions of the ER highlighted in yellow. **F)** Quantification of LD diameter in wild-type and *pex30*Δ cells. n= 50 cells. **G,H)** Cells with grown to early stationary growth phase in SC medium containing [^3^H]acetate and the amount of TAG (I) and SE (J) determined. **I)** Cells expressing Dga1-GFP from a plasmid were grown to stationary growth phase in SC (0 hours). Cells were washed, resuspended in fresh media, and the percent with Dga1-GFP only on LDs over time determined. Bottom panels show examples of Dga1-GFP distribution patterns: ER only (left), ER and LDs (middle), and LD only (right). G,H, and I show mean ± s.d. of three independent experiments, * p < 0.05, ** p< 0.005, *** p< 0.0005, student’s t-test. Bars = 3μm.

Four proteins in yeast contain reticulon homology domains (RHDs) like that of Pex30: Pex28, Pex29, Pex31, and Pex32. We determined whether mutants lacking these proteins had changes in LD size, LD clustering, or neutral lipids levels but found they did not, though there was some LD clustering in cells lacking Pex29 (Fig. 2G,H and Supplementary Fig. 2). These findings suggest the functions of Pex30 do not overlap with those of similar RHD-containing proteins.

Since LDs frequently associate with Pex30-containing portions of the ER, we wondered whether ER-LD contacts were altered in cells lacking Pex30. In yeast, LDs remain attached to the ER by a membrane neck that is thought to be formed from the cytoplasmic leaflet of the ER membrane and is continuous with the phospholipid monolayer surrounding LDs^12^. Dga1 and some other ER proteins with hairpin-like membrane domains can diffuse between the ER and LDs^12^^,^^23^^,^^24^. Thus, if ER-LD contacts are altered in *pex30*Δ cells, Dga1 movement between these organelles might be affected. We found that Dga1-GFP is primarily on the ER in late stationary growth phase but diffuses onto LDs when cells are transferred to fresh media. We determined the percent of cells that have Dga1-GFP only on LDs after cells were shifted to fresh media. Two hours after shift, there were significantly fewer *pex30*Δ than wild-type cells with Dga1-GFP on LDs (Fig. 2I). Interestingly, after 24 hours, when cells have returned to stationary growth phase, more *pex30*Δ cells than wild-type cells still have Dga1-GFP on LDs (Fig. 2I). These results suggest that Dga1-GFP movement between the ER and LDs slows in cell lacking Pex30, consistent with the idea that ER-LD contacts are altered in cells lacking Pex30.

Since seipin has been suggested to localize at sites where LDs are associated with ER^2^^,^^25^, we wondered how LD biogenesis would be affected in cell lacking both Pex30 and seipin (Sei1). Surprisingly, these cells (*sei1pex30*Δ) have a substantial growth defect (Fig. 3A). The defect was corrected when only the RHD-containing N-terminal 234 residues of Pex30 were expressed in the *sei1pex30*Δ cells, indicating that the membrane-shaping function of Pex30 is necessary to support optimal growth of cell lacking seipin (Fig. 3A). Interestingly, elimination of seipin causes a profound redistribution of Pex30 in the ER; Pex30-2xmCherry accumulates in a single punctum that localizes with the LD marker Nem1-GFP (Fig. 3B), indicating that seipin affects the distribution of Pex30 subdomains.

**Figure 3.**
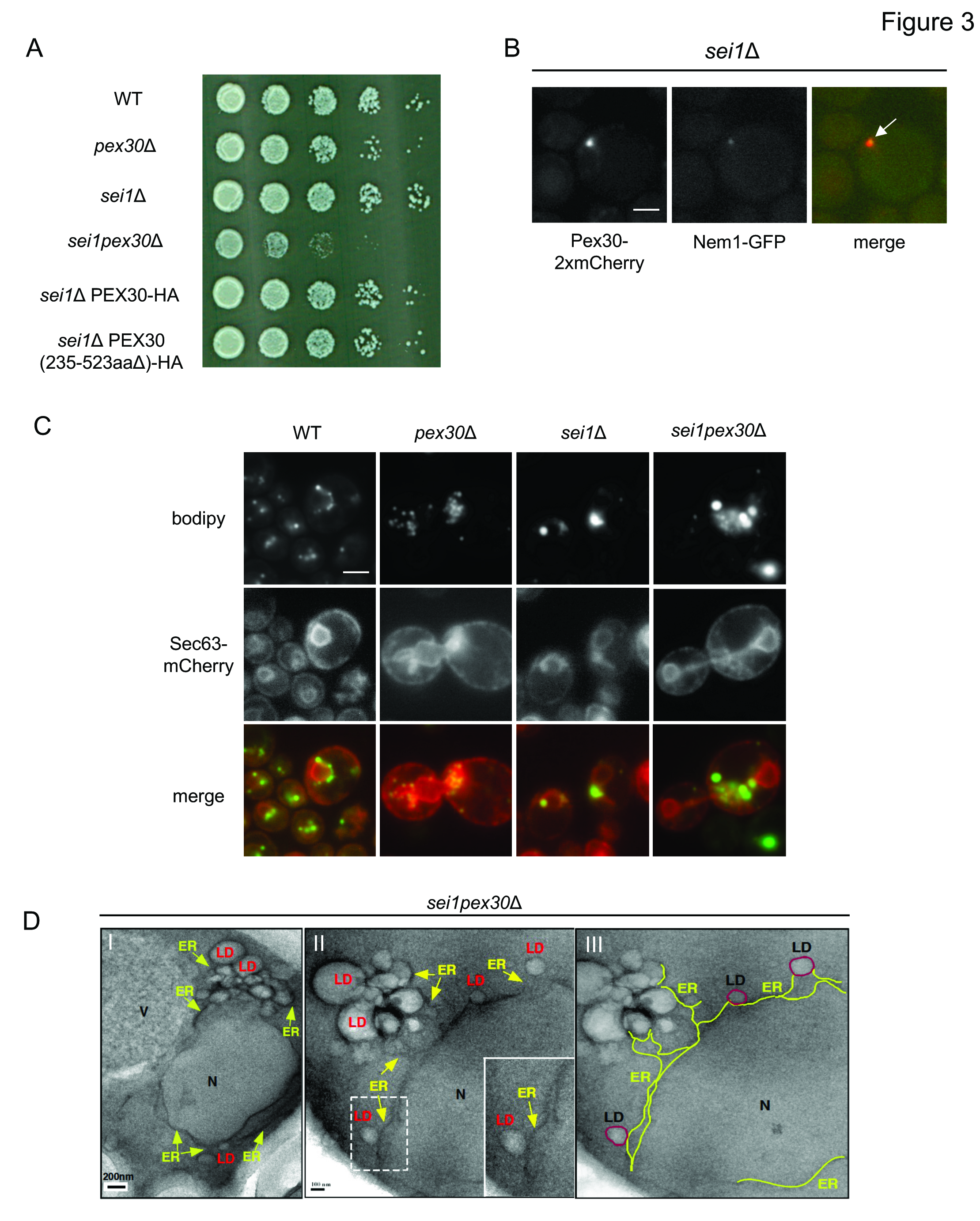
Cells lacking Pex30 and seipin have altered LDs and growth defects. **A)** Strains were grown to mid-logarithmic growth phase, serially diluted, spotted on to the YPD plates and incubated at 23°C for 3 days. **B)** A *sei1*Δ mutant expressing endogenously tagged Pex30-2xmCherry and Nem1-GFP was visualized live. Arrow indicates colocalization. Bar = 3μm. **C)** Images of cells stained with BODIPY and expressing Sec63-mCherry from a plasmid. Bar = 3μm. **D)** Cells growing in SC without oleic acid were fixed and visualized by EM, abbreviations as in Fig. 2B-E. III shows the same image as II but with portions of the ER highlighted in yellow.

It is not clear why *sei1pex30*Δ cells have a growth defect. We found that *sei1pex30*Δ cells form large clusters of small and big LDs and the ER associated with the LDs is highly proliferated around the LDs (Fig. 3C and 3D). These changes could affect ER function and cause a growth defect. Alternatively, Pex30 and seipin may share a common unknown function, perhaps related to lipid metabolism. Together, these findings provide additional evidence that Pex30 plays an important role in LD biogenesis and function and suggest that Pex30 and seipin may have partially overlapping functions.

Pex30 does not have a mammalian homologue but we wondered whether there is an RHD-containing protein in higher eukaryotes that plays a similar role. Using the structural homology prediction program HHpred^26^, we found that the human protein MCTP2 has a RHD similar to that of Pex30 (Fig. 4A). MCTP2 is an 878 amino acid ER resident protein with a membrane embedded regions containing the RHD near the C-terminus and three C2 domains ^27^^,^^28^. MCTP proteins are conserved in higher eukaryotes. *Drosophila* and *C. elegans* have one MCTP whereas humans contain MCTP1 and MCTP2^27^. Here we show that human MCTP2 is a functional homologue of Pex30. We expressed the C-terminal 237 amino acids of MCTP2, which contain the RHD, fused to YFP (YFP-MCTP2) in *S. cerevisiae* under the strong *RTN1* promoter. YFP-MCTP2 complements the growth defect of *sei1pex30*Δ cells (Fig. 4B). Cells lacking the reticulons, Rtn1 and Rtn2, and the reticulon-like protein Yop1 (*rtn1rtn2yop1Δ*), have a defect in ER structure^29^ that is corrected by overexpression of Pex30^5^. We found that YFP-MCTP2 similarly restores ER structure. The cortical ER forms large sheet-like structures in *rtn1rtn2yop1*Δ cells that are not present in wild-type cells, which contains largely tubular ER in the cortex. We found that cortical ER structure in *rtn1rtn2yop1*Δ cells becomes tubular when YFP-MCTP2 is expressed in these cells (Fig. 4C). We previously found that *rtn1rtn2yop1*Δ cells that also lack the lipid regulator Spo7 are not viable but grow when Pex30 is overexpressed^5^. Similarly, overexpression of YFP-MCTP2 also rescued the *rtn1rtn2yop1spo7*Δ mutant (Supplementary Fig. 3A). Together, these findings indicate that YFP-MCTP2 is an ER-shaping protein that can functionally replace Pex30 in yeast.

**Figure 4.**
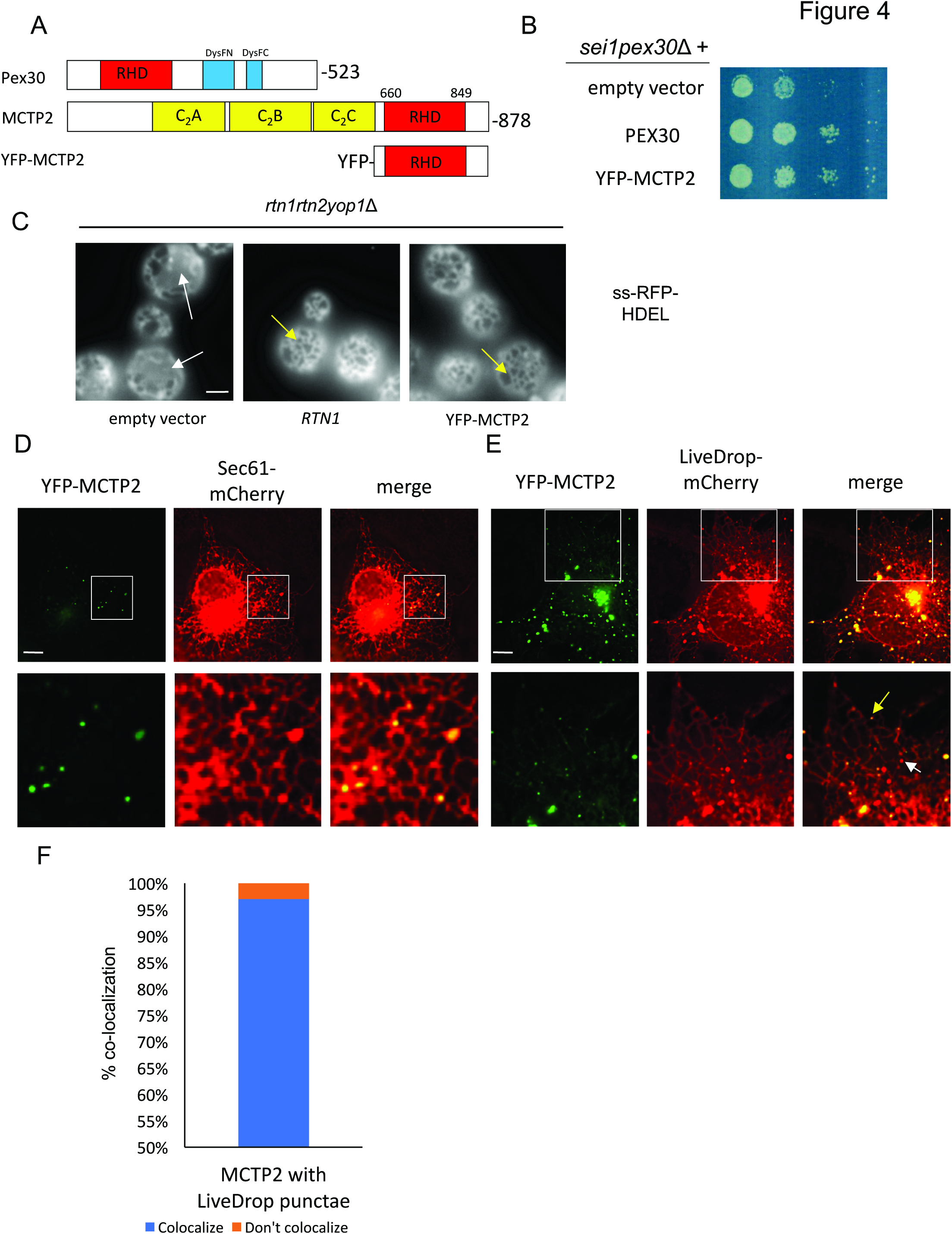
Human MCTP2 is a functional homologue of Pex30. **A)** Domains in Pex30, MCTP2, and YFP-MCTP2. MCTP2 has 3 C2 domains A-C. Pex30 contains a dysferlin (DysF) domain that is divided into N and C portions. **B)** Strains were grown to mid-logarithmic growth phase, serially diluted, spotted on to SC plates and incubated at 23^o^C for 3 days. **C)** Images of the *rtn1rtn2yop1*Δ cells expressing the ER marker ss-RFP-HDEL and Rtn1or YFP-MCTP2. Images are of the cell periphery. White arrows indicate ER sheets and yellow arrow show regions containing ER tubules. Bar = 3μm. **D,E)** Images of COS7 cells co-transfected with plasmids expressing YFP-MCTP2 and Sec61-mCherry (D) or LiveDrop-mCherry (E). Yellow arrow indicates colocalization and white arrow indicates a LiveDrop puncta that does not colocalize with YFP-MCTP2. Stacks of 20 images with a step size of 0.3μm were taken and deconvolved; images from single plane are shown. Bars = 5 μm. **F)** Quantification of colocalization of punctae from (E). n =300 punctae.

We found that YFP-MCTP2 localizes to ER subdomains in mammalian cells. YFP-MCTP2 was transiently expressed in COS7 cells together with the ER marker Sec61-mCherry. When YFP-MCTP2 was expressed at low levels, it was found in puncta in the ER (Fig. 4D). These puncta are stable and remain associated with the same region of the ER over time (Supplementary video 1). When expressed at high levels, YFP-MCTP2 localized to ER tubules and the edges of ER sheets (Supplementary Fig. 3B), a localization shared with the reticulons^30^ and consistent with the idea that the C-terminal region of MCTP2 contains a RHD.

We next determined whether MCTP2-subdomains are sites of LD biogenesis and associate with LDs. COS7 cells were transiently transfected with YFP-MCTP2 and LiveDrop-mCherry, a fusion protein demonstrated to target nascent LDs forming in the ER and mature LDs^3^. We found that most MCTP2 and LiveDrop-mCherry punctae colocalized (Fig. 4E and 4F). LiveDrop-mCherry punctae that did not colocalize with YFP-MCTP2 were largely not associated with the ER (Fig. 4E). These findings indicate that YFP-MCTP2 localizes to ER sites where new LDs form and suggest that ER domains containing YFP-MCTP2 are often associated with mature LDs. Thus, MCTP2 may play a role in LD biogenesis in mammalian cells like that of Pex30 in yeast.

We wondered whether Pex30/MCTP2 sites in the ER not only associate with LDs but also with PPVs and peroxisomes, since we previously found that PPVs, are generated at the Pex30 subdomain^5^. To visualize PPVs in *S. cerevisiae*, we used strains that lack the proteins Pex3 and Atg1 (*pex3atg1Δ*). Pex3 is required for peroxisome biogenesis and it had been thought that cells lacking this protein do not contain PPVs^31^. However, it was subsequently discovered that PPVs are present in *pex3Δ* cells when they also lack Atg1, which is required for autophagy^32^. These PPVs contain Pex14-GFP and there are typically 1 or 2 PPVs per cell. We found that some Pex14-GFP puncta are on vesicles while others are on the ER, at Pex30 subdomains^5^. To colocalize PPVs, LDs and Pex30 subdomains, we expressed Pex14-GFP, Pex30-2xmCherry, and the LD marker Erg6-BFP in *pex3atg1Δ* cells. Remarkably, most PPVs are closely associated or colocalized with Pex30 subdomains and LDs (Fig 5A and Supplementary Fig. 4). The association between PPVs and LDs is even more pronounced in *pex3atg1Δ* cells that also lack seipin (Supplementary Fig. 4). These findings reveal that PPVs and LDs often remain associated with the same Pex30 subdomain and suggest that a single subdomain may generate both PPVs and LDs. It is also possible that both PPVs and LDs form at different regions in the ER but subsequently associate with the same Pex30 subdomain.

Pex30 subdomains are also associated with mature peroxisomes in yeast^33^. We found that MCTP2 subdomains similarly associate with both LDs and mature peroxisomes in mammalian cells. YFP-MCTP2, LiveDrop-mCherry, and the peroxisome marker CFP-SKL were co-expressed in COS7 cells. LDs and peroxisomes often associate with the same MCTP2 subdomain (Fig. 5B). About 30 percent of the MCTP2 and LiveDrop punctae that colocalize are also associated with peroxisomes (Fig. 5C). These peroxisomes are either transiently (Supplementary video 2A) or stably (Supplementary video 2B) associated with the ER subdomains containing MCTP2 and LiveDrop.

Collectively, our findings indicate that conserved ER subdomains with specialized RHD-containing proteins are sites where new LDs are formed. These sites are likely the location in the ER where LD formation begins since they are enriched in the TAG precursor DAG. Mature LDs often remain associated with these ER subdomains. We show that Pex30/MCTP2 subdomains are sites of biogenesis of both LDs and peroxisomes and thus may coordinate the formation of these organelles in response to changes in lipid metabolism. Interestingly, individual Pex30/MCTP2 subdomains often remain associated with both mature LDs and peroxisomes. Since, LDs and peroxisomes are also known to make close contacts^34^^,^^35^, Pex30/MCTP2 subdomains may facilitate intracellular signaling between the ER, peroxisomes, and LDs.

**Figure 5.**
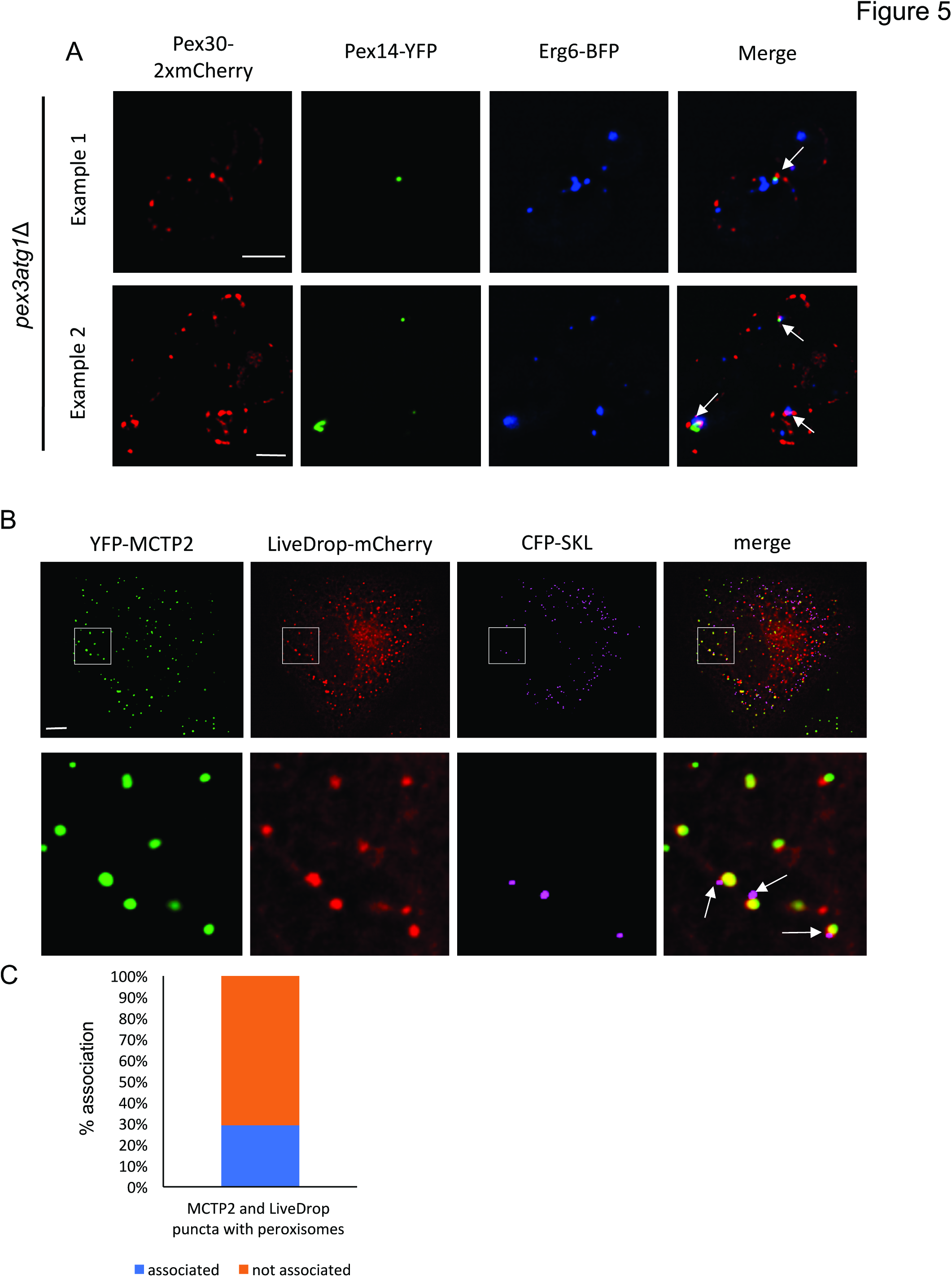
LD and peroxisome biogenesis occurs at the same ER subdomains. **A)** Images of *pex3atg1*Δ cells expressing endogenously tagged Pex30-2xmCherry, Pex14-YFP (a PPV maker), and Erg6-BFP (an LD marker) from plasmids. Stacks of 3 images with a step size of 0.25μm were taken and deconvolved; images from single plane are shown. Bar = 3μm. **B)** Images of COS7 cells transiently co-transfected with plasmids expressing YFP-MCTP2, LiveDrop-mCherry (an LD marker), CFP-SKL (a peroxisome marker). Stacks of 20 images with a step size of 0.3μm were taken and deconvolved; images from single plane are shown. Bar = 5 μm. **C)** Quantification of percent colocalization of punctae from (B), n =300 punctae.

## AUTHOR CONTRIBUTIONS

ASJ and WAP designed the experiments. ASJ and VC performed the experiments. TPL performed the HHpred analysis. ASJ and WAP analyzed the data and wrote the manuscript.

## ACKNOWLEDGEMENTS

This work was supported by the Intramural Research Program of the National Institute of Diabetes and Digestive and Kidney Diseases. TPL was supported by the Biotechnology and Biological Sciences Research Council Bioinformatics and Biological Resources Fund (grant BB/M011801). We thank T. Rapoport, T. Sudof, T. Walther. T. Balla, and J. Lippincott-Schwartz for providing reagents, J. Cooper for use of her Deltavision microscope and T. Balla and Y. Ye for critically reading the manuscript.

## METHODS

### Yeast strains and plasmids

The strains and plasmids used in this study are listed in Tables S1 and S2. Deletion strains were constructed by mating or PCR-based targeted homologous recombination to replace the ORF of genes of interest with cassettes (Longtine et al., 1998). PCR-based targeted homologous recombination was also used to generate endogenously expressed C-terminally tagged fusion proteins. The 2xmCherry-*URA3* and GFP-*HIS3MX6* cassettes were obtained from O. Cohen-Fix (National Institutes of Health/National Institute of Diabetes and Digestive and Kidney Diseases, Bethesda, MD), YFP-*KanMX6* from J. Cooper (National Institutes of Health/National Cancer Institute, Bethesda, MD), and yEmCherry-*HIS5MX6* and pRS305-*PHO8*-3xBFP from J. Nunnari (University of California, Davis, Davis, CA).

The plasmid encoding ER-DAG sensor was constructed by fusing the portion of the human protein kinase D gene encoding amino acids 136-343 (obtained from Tamas Balla, National Institute of Child Health and Human Development, NIH) to genes encoding GFP and the tail-anchored transmembrane domain of *S. cerevisiae* Ubc6 under the *ADH1* promoter in the plasmid YEplac181. The plasmids used in live cell imaging of COS7 cells were Sec61-mCherry from T. Rapoport (Harvard University), YFP-MCTP2 from T. Sudof (Stanford University), LiveDrop-mCherry from T. Walther (Harvard University) and CFP-SKL from J. Lippincott-Schwartz (Janelia Research).

### Media and growth conditions

Yeast cells were grown at 30°C, unless otherwise indicated, in YPD medium (1% Bacto yeast extract, 2% Bacto Peptone, and 2% glucose) or in synthetic complete (SC) media containing 2% glucose, 0.67% yeast nitrogen base without amino acids (United States Biological) and an amino acid mix (United States Biological). In some cases the glucose in SC was replaced with 2% raffinose or 2% galactose. When inducing LD production, cells were washed with sterile water twice and transferred to SC containing 1mM oleic acid and 1% Brij58. Images were taken within 1-2 hours. When staining LDs with BODIPY, cells in early stationary growth phase were washed with phosphate buffered saline and incubated with 0.5μg/ml BODIPY 493/503 (Invitrogen) for 10 minutes.

COS7 cells were cultured in DMEM (Gibco) supplemented with 10% FBS (Gibco) and 2 mM L-glutamine (Gibco) at 37°C in humidified air containing 5% CO_2_. Prior to live cell imaging, the medium was changes to CO_2_-independent medium (Gibco) containing 10% FBS and 2 mM L-glutamine.

### Fluorescence microscopy

For Fig. 2A, 2I, 3B, 3C, 4C, Supplementary fig. 2H and supplementary Fig. 4, cells were imaged live in growth media using a BX61 microscope (Olympus) with a UPlanAPO Å~100/1.35 lens and a Retiga EX camera (QImaging) and processed using iVision software (version 4.0.5). For Fig. 1, 4D, 4E, 5A, 5B, Supplementary Fig. 1 and Supplementary Fig. 3, imaging was performed at 30°C in an Environmental Chamber with a DeltaVision Spectris (Applied Precision Ltd.) comprising a wide-field inverted epifluorescence microscope (IX70; Olympus), a 100Å~ NA 1.4 oil immersion objective (UPlanSAPO; Olympus), and a charge-coupled device Cool-Snap HQ camera (Photometrics). For COS7 cells imaging, cells were cultured in MatTek 35mm petri dish, 14mm microwell, No. 1.5 coverglass, (0.16mm-0.19mm). Cells were transfected with indicated plasmids using Lipofectamine 2000 (Invitrogen) according to the manufacturer’s instructions. Time-lapse images were acquired every 60 seconds for 10 minutes for Supplementary Fig. 2 and every 12 seconds for 2.5 minutes for Supplementary Fig. 3. Images were deconvolved (conserved ratio method) using SoftWorx (Applied Precision Ltd.). Brightness and contrast were adjusted using Photoshop CC (Adobe Systems).

### Purification of MBP-Pex30 (250-523) protein

The portion of *PEX30* encoding amino acids 250-523 was cloned into the plasmid pMAL-c2x. The plasmid was expressed *E.coli* grow to mid-logarithmic growth phase at 37°C, 0.2mM IPTG was added to the medium, and the cell were grown at 30°C for 3 hours. The cells were harvested by centrifugation, resuspended in lysis buffer (200mM NaCl, 20mM TRIS pH 8.0, 1mM EDTA, 1mM DTT and protease inhibitor), and lysed using french press. The soluble protein was purified from the lysate using amylose resin (New England Biolabs) on an FPLC column (GE healthcare). Protein was eluted using 10mM maltose. The purified protein was separated using HiLoad 16/60 Superdex 200 prep grade (GE healthcare). Peak fractions with purified protein were collected, pooled and analyzed using SDS-PAGE stained with Coomassie. Purified protein (0.5μg/ml) was used for *in vitro* lipid binding assay.

### Electron microscopy (EM)

Sample preparation and visualization was performed as described previously^6^. In brief, yeast cells were grown to mid-logarithmic growth phase, and 10 OD_600_ units of cells were pelleted and fixed in 1 ml of fixative media (2.5% glutaraldehyde, 1.25% PFA, and 40 mM potassium phosphate, pH 7.0) for 20 min at room temperature. Cells were pelleted, resuspended in 1 ml fresh fixative media, and incubated on ice for 1 h. The cells were pelleted, washed twice with 0.9% NaCl, once with water, incubated with 2% KMnO_4_ for 5 min at room temperature, centrifuged, and resuspended in 2% KMmO_4_ for 45 min at room temperature for en bloc staining. The cells were dehydrated using ethanol, embedded using Spurr’s resin (Electron Microscopy Sciences), and polymerized as described previously. Semi-and ultrathin sections were produced with a diamond knife (DiATO ME) on an ultra-microtome (Ultracut UCT; Leica Biosystems), collected on 200 mesh copper grids (Electron Microscopy Sciences), poststained with uranyl acetate and lead citrate, and visualized with a Tecnai T12 transmission electron microscope (FEI), operating at 120 kV. Pictures were recorded on a bottom-mounted 2k Å~ 2k CCD camera (Gatan). Brightness and contrast were adjusted to the entire images using Photoshop (version CC 2014).

### Neutral Lipids detection

Cells were grown to early stationary growth phase in SC glucose containing 10 μCi/ml [^3^H] acetate (American Radiolabeled Chemicals). 10 OD_600_ units of cells were harvested, lysed using a Precellys24 homogenizer, and lipids were extracted as described previously^36^. To quantitate TAG and SE, the lipids were spotted onto silica gel 60 TLC plates (EMD Millipore) and developed with hexane-diethylether-acetic acid (80:20:1). Lipids on TLC plates were quantified with a RITA Star Thin Layer Analyzer (Raytest).

### DGA1-GFP localization assay

Cells expressing Dga1-GFP from the *DGA1* promoter in the centromeric plasmid YCplac111 were grown in SC. These cells were diluted in fresh medium and incubated for 24 hours followed by dilution to 0.3 OD_600_ units/ml. These cells were then imaged at the indicated times to determine the localization of Dga1-GFP.

---

**Supplement figure 1.**
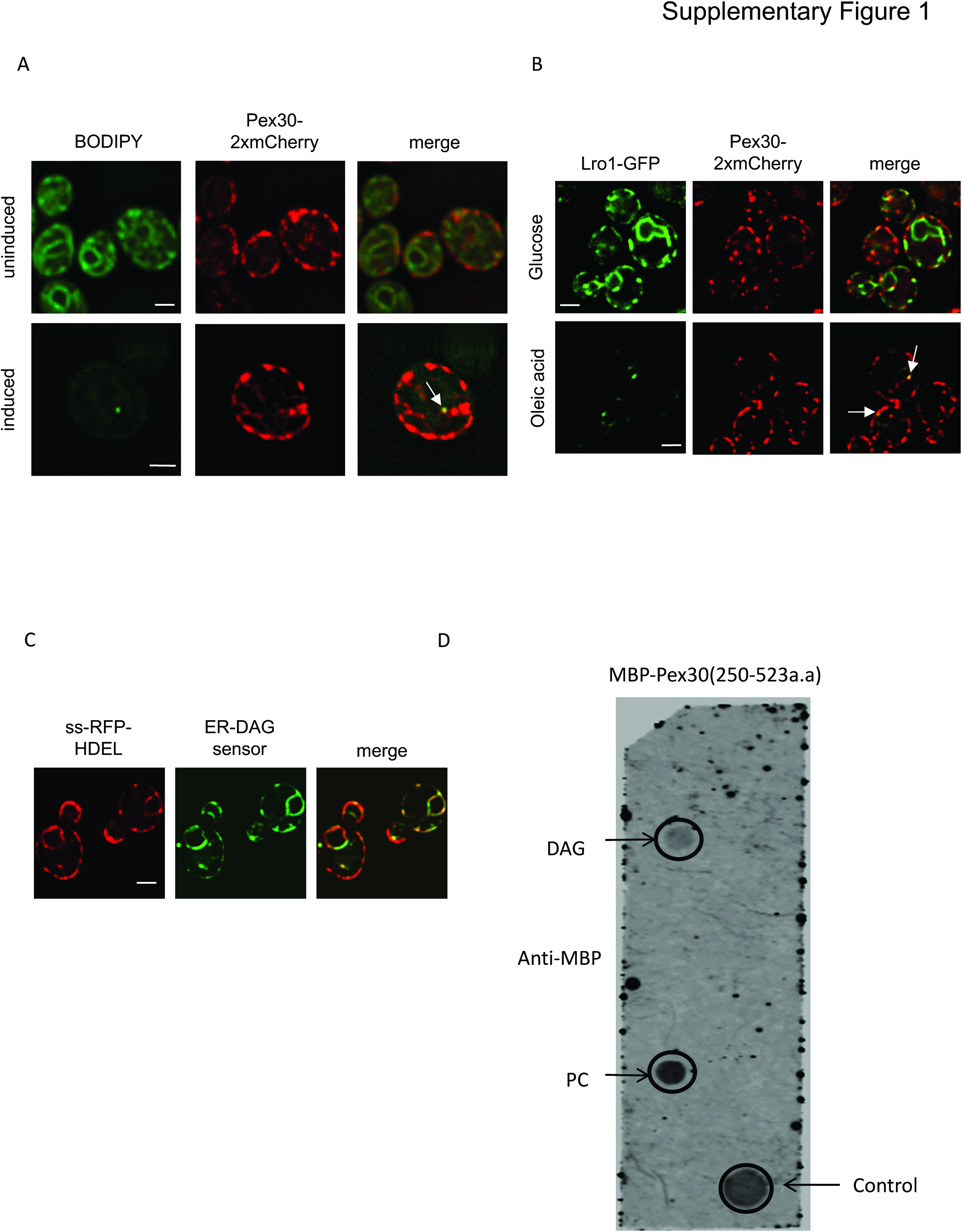
Data associated with Fig. 1. **A)** Same as Fig. 1A except that LDs are visualized using BODIPY instead of Erg6-GFP. Bar = 3μm. **B)** Same as Fig. 1C except cells are expressing endogenously tagged Pex30-2xmCherry and Lro1-GFP from a plasmid. Bar = 3μm. **C)** Images of cells expressing endogenously tagged ss-RFP-HDEL (an ER maker) and ER-DAG sensor from a plasmid. **D)** The C-terminal 273 amino acids of Pex30 fused to MBP were purified from *E.coli*. The purified was protein (0.5μg/ml) was incubated with a lipid strip (Echelon Inc., catalog # P-6002) for 1 hour at room temperature. The lipid strip was immunoblotted with anti-MBP antibody.

**Supplement figure 2.**
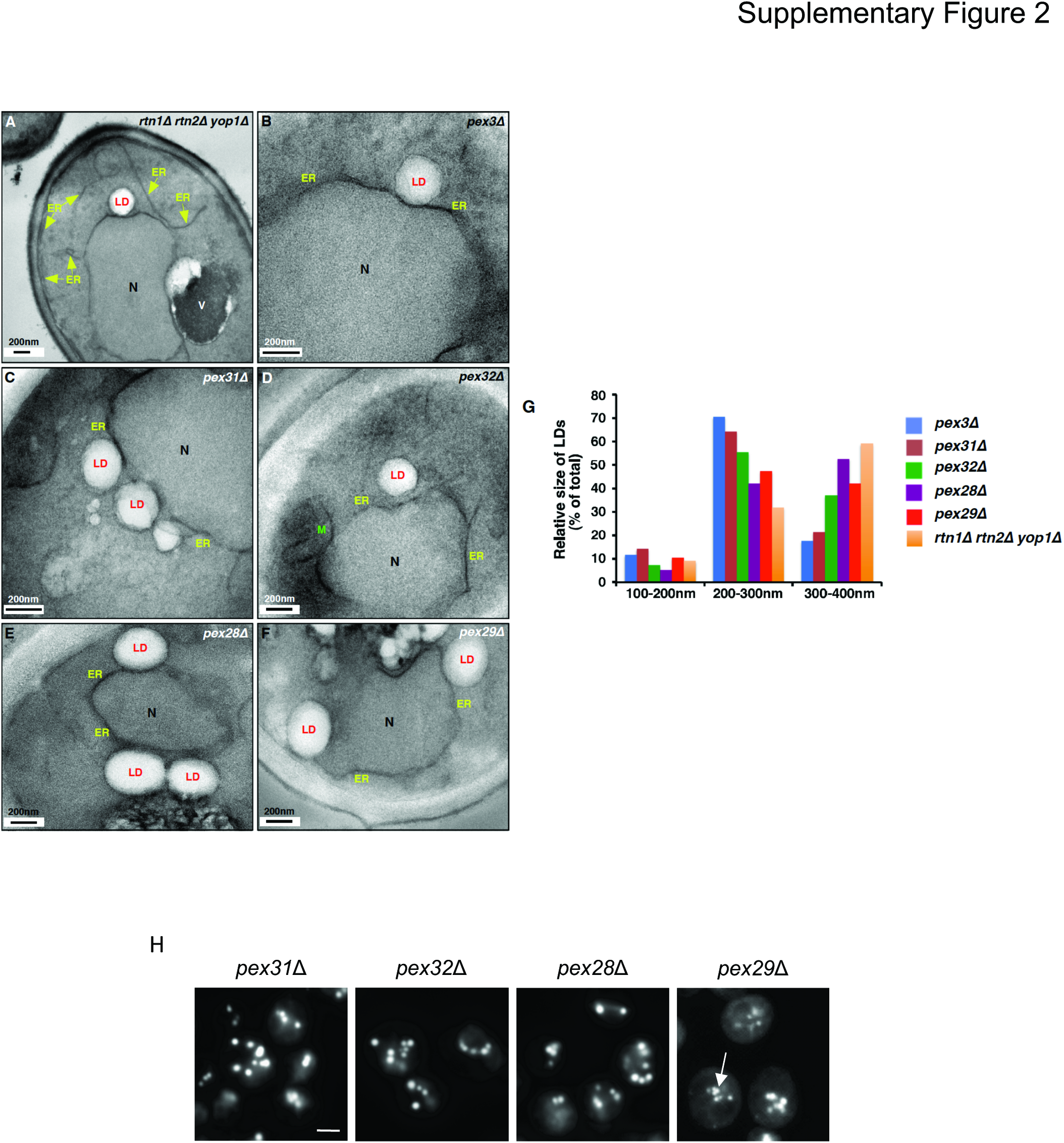
Data associated with Fig. 2. **A-F)** Cells growing in SC were fixed and visualized by EM. Yellow arrows indicate ER. V; vacuole, N; Nucleus, LD; Lipid droplets. E shows the same image as D but with portions of the ER highlighted in yellow. **G)** Quantification of LD diameter in wild-type and *pex30*Δ cells. n= 20 cells. **H)** Cells were grown to early stationary growth phase in SC and stained with BODIPY to visualize LDs. Stacks of 10 images with a step size of 0.2μm were taken; images from a single plane are shown. Arrows indicate clustered LDs. Bar = 3μm.

**Supplement figure 3.**
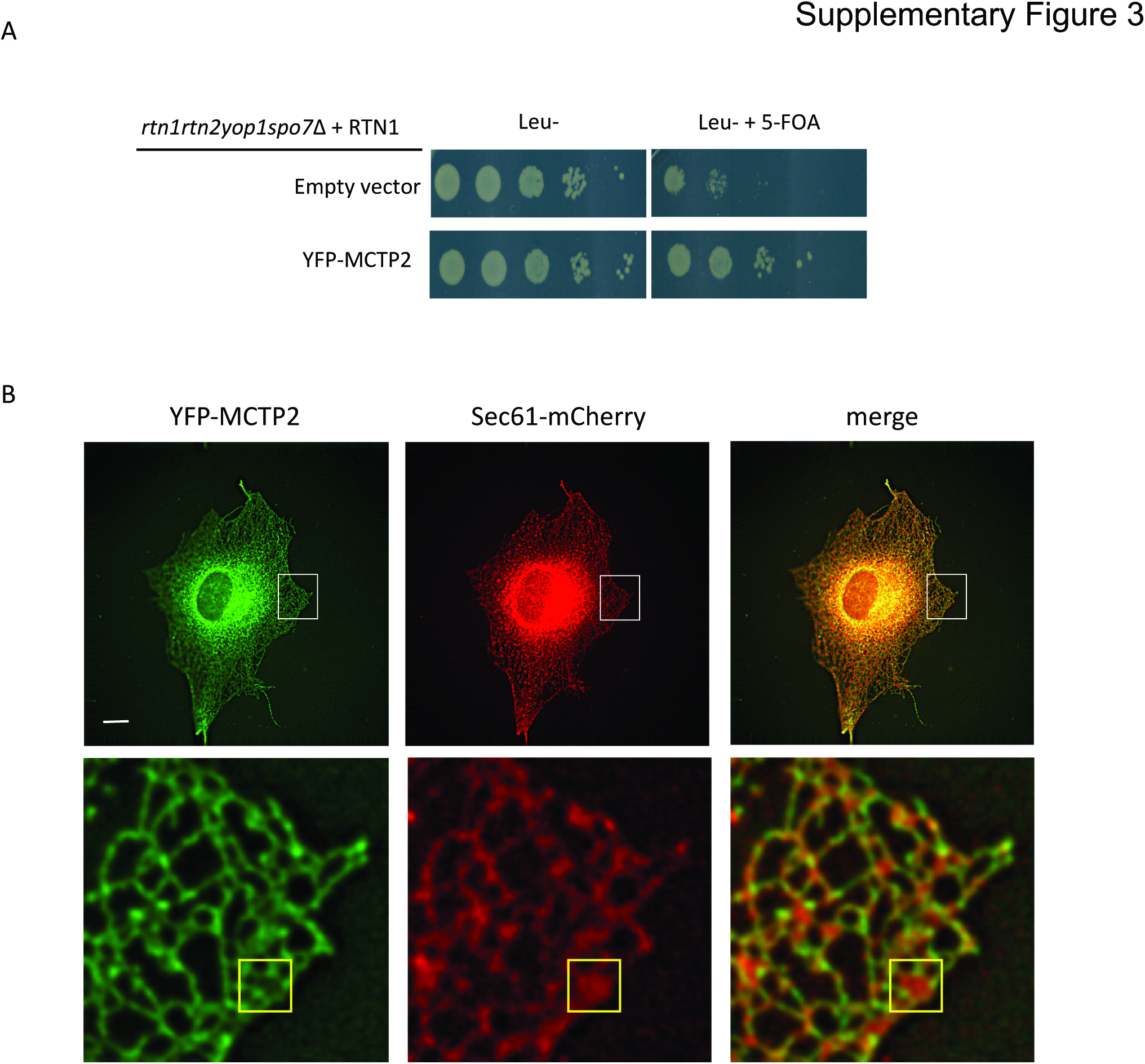
Data associated with Fig. 4. **A)** Cells lacking Rtn1, Rtn2, Yop1, and Spo7 (*rtn1rtn2yop1spo7*Δ) are not viable unless they contain a plasmid expressing an RHD-containing protein. The *rtn1rtn2yop1spo7*Δ cells were complemented with a plasmid containing *RTN1* and *URA3*, which can be counter selected with 5-Fluoroorotic Acid (5-FOA). This strain containing and empty vector or a plasmid expressing YFP-MCTP2 were grown to mid-logarithmic growth phase, serially diluted, spotted on to SC plates with or without 5-FOA, and incubated at 30°C for 3 days. **B)** Same as Fig.4D but showing a cell expressing YFP-MCTP2 at a high level. Region in white box shown in higher magnification in lower panels. Yellow box shows ER sheet with YFP-MCTP2 at edges. Bar = 5μm.

**Supplement figure 4.**
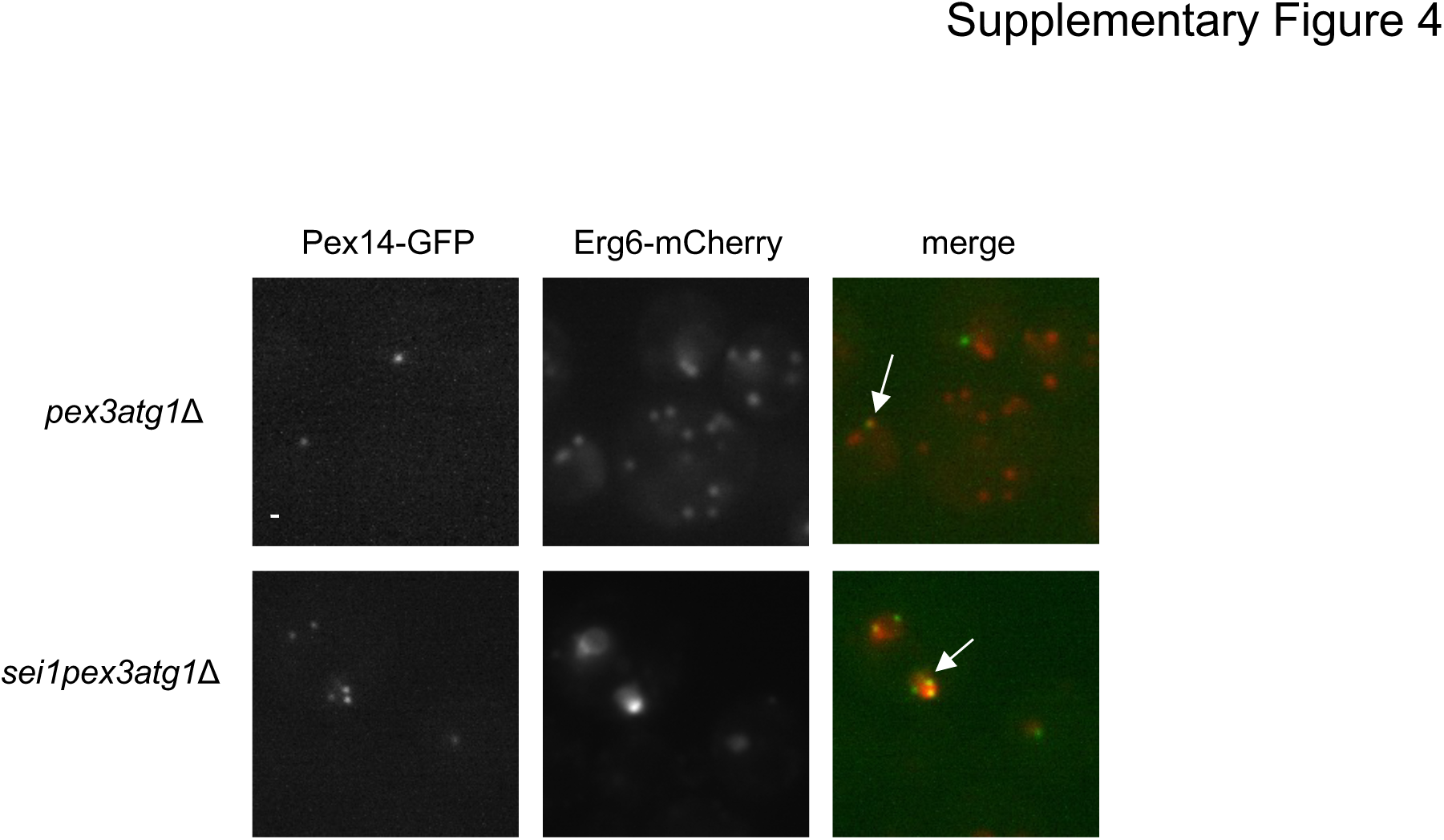
Data associated with Fig. 5. PPVs are closely associated with LDs in *pex3atg1*Δ cells. Images of cells expressing endogenously tagged Pex14-GFP and Erg6-mCherry. Stacks of 10 images with a step size of 0.2μm were taken; images from a single plane are shown. White arrows indicate co-localization. Bar = 3μm.

**Supplemental video 1. YFP-MCTP2 is stably associated with ER subdomains.**

Time-lapse images of COS7 cells co-transfected with plasmids expressing Sec61-mCherry and YFP-MCTP2. Stacks of 20 images with a step size of 0.3μm were taken for 10 minutes with time interval of 60 seconds and deconvolved; images from single plane are shown.

**Supplemental video 2: Peroxisomes associate with ER subdomains containing YFP-MCTP2 and LiveDrop-mCherry.**

Peroxisomes are either transiently (A) (one punctae in the center) or stably (B) (two puncta in the center) associated with ER subdomains containing MCTP2 protein. Time-lapse images of COS7 cells co-transfected with plasmids expressing Sec61-mCherry, CFP-SKL and YFP-MCTP2. Stacks of 20 images with a step size of 0.3μm were taken for 2.5 minutes with time interval of 12 seconds and deconvolved; images from single plane are shown.

